# Organohalide respiration by a *Desulforhopalus*-dominated community

**DOI:** 10.1101/2023.04.18.537297

**Authors:** Chen Zhang, Siavash Atashgahi, Tom N.P. Bosma, Hauke Smidt

## Abstract

Despite the fact that several potential organohalide-respiring bacteria (OHRB) were discovered in metagenome-assembled genomes (MAGs) in our previous study of marine sediments from Aarhus Bay, delineation of their roles and interactions are yet to be disentangled. Henceforth, obtaining corresponding pure cultures or more defined consortia would be highly instrumental for more detailed eco-physiological studies. To this end, we isolated a colony from an anaerobic slant tube culture inoculated with a stable PCE dehalogenating enrichment. Intriguingly, the derived culture exhibited debromination only, instead of PCE dechlorination, under sulfate-reducing conditions. The culture was capable of conserving energy for growth via debromination of 2,6-dibromophenol (2,6-DBP). Analysis of 16S rRNA gene sequence data extracted from shot gun metagenome sequences revealed that a strain belonging to *Desulforhopalus* was the predominant member of the consortium at a relative abundance of 29 %. Moreover, five bins (completeness > 85% and contamination < 3%) were assembled and all were identified as potentially new species (average nucleotide identity, ANI < 95%). Two bins from potential OHRB, bin.3 belonging to *Desulfoplanes*, and bin.4 belonging to *Marinifilaceae*, were found to encode reductive dehalogenase (RDase) genes, whereas bin.5 was found to contain a gene coding for thiolytic tetrachloro-p-hydroquinone (TPh-) RDase bearing 23.4 % identity to TPh-RDase of *Sphingobium chlorophenolicum*. The expression of all three RDase genes was strongly-induced after adding 2,6-DBP. Acetylene, a known inhibitor of different redox-active metalloenzymes, was found to inhibit methanogenesis as well as reductive dehalogenation without affecting gene expression, suggesting post-transcriptional inhibition. Phylogenomic analyses revealed the ecological importance of complementary roles of community members, including complete *de novo* vitamin B12 biosynthesis, which agreed with physiological data. Altogether, the findings presented here provided insight into the mutualism of the consortium and provided leads for synthetic OHR community optimization strategies for *in situ* bioremediation.

## Introduction

Owing to recent technological and conceptual advances particularly regarding DNA and RNA sequence based approaches, an ever increasing number of studies is being conducted where such approaches provide substantial support for understanding the biological activity in underexplored biospheres, also expanding our knowledge on evolutionary aspects across the tree of life (1–3). Recently, accumulating evidence revealed the presence of RDase genes in marine sediments of Aarhus Bay, and detection of corresponding transcripts suggested the occurrence of organohalide respiration (OHR), further diversifying the spectrum of local energy-conserving mechanisms, especially under low nutrient conditions such as those prevailing in marine sediments (4, 5). In addition, a high Br^-^/Cl^-^ ratio alongside the depth of sediments suggested the prevalence of reductive debromination (5). On this basis, experiments described in previous work eco-physiologically validated the OHR potential of Aarhus Bay sediments on tetrachloroethane (PCE), 2,6-dibromophenol (2,6-DBP), 3-bromophenol (3-BP), 1,4-dibromobenzene (1,4-DBB), and 2,4,6-triiodophenol (2,4,6-TIP) (6). Subsequent metagenomic and -transcriptomic analyses allowed for the assembly of metagenome assembled genomes (MAGs), representing a diverse group of novel, potential organohalide respiring bacteria (OHRB), most of which showed induction of the expression of genes related to OHR in the presence of PCE (7).

Reductive dehalogenation is catalyzed by reductive dehalogenases (RDases), which in general are classified into two types due to their differences in catalytic mechanism: 1) RDases in anaerobes, and particularly OHRB (8, 9); 2) thiolytic tetrachloro-p-hydroquinone (Tph)-RDase in aerobes (10–12). Both enzymes require anoxic conditions for their catalytic activity (13, 14). Beyond their role in the biotransformation of organohalides, respiratory RDases have been shown to be coupled to energy conservation for bacterial growth, which is not the case for TPh-RDase. Therefore, respiratory RDases have attracted more research interests. In spite of sequence dissimilarities, several functional motifs of respiratory RDases are conserved, including a twin-arginine translocation (Tat domain, RR) signal peptide, and two Iron-Sulfur cluster motifs (Fe-S1: FCXXCXXCXXXCP and Fe-S2: CXXCXXXCP) involved in the electron transfer from the electron donor to the catalytic site of the enzyme (15–18). In addition, the operon that encodes the respiratory RDase, also named RdhA or RDase A, in most cases also encodes a second protein, RDase B, that acts as an anchor protein to localize RDase A to the outer face of the cytoplasmic membrane (17). TPh-RDases were classified into the Glutathione-S-Transferase (GST) superfamily, in which cysteine and serine are two conserved catalytic site residues that are critical for the catalytic activity of Tph-RDases that use glutathione as the reducing equivalent (19). Interestingly, our previous data revealed the presence of a TPh-RDase gene in bin.15, which was affiliated with the strictly anaerobic genus *Desulforhopalus* (7, 20, 21). Moreover, the TPh-RDase gene was also induced during PCE dehalogenation regardless of the additional sulfate. Nevertheless, a role of TPh-RDase under anoxic conditions and its relationship with the respiratory RDases found in other MAGs awaits further study.

The activity of respiratory RDases requires a corrinoid cofactor. Previous work on *Dehalococcoides* has shown that they lack *de novo* corrinoid biosynthesis capacity, and that absence of corrinoids abolished RDase activity (22–25). Thus, members of *Dehalococcoides* require the supplementation of external corrinoids for reductive dehalogenation (22). With more OHRB isolated and characterized, few OHRB, such as *Sulfurospirillum multivorans*, have been shown to not only utilize a range of different electron donors and carbon sources, but also self-supply the necessary corrinoid cofactor via a complete biosynthesis pathway (26). A recent coculture study of *D. mccartyi* strain 195 and *S. multivorans* achieved the complete PCE dehalogenation to ethene over three times faster as compared to a monoculture of *D. mccartyi* strain 195 without adding corrinoids (27). This was the first synthetic coculture establishment, which successfully showed the conceptual feasibility of synthesizing an OHR community, however, following a bottom-up approach for establishing defined synthetic consortia remains challenging due to the complex ecological and physiological interactions among the members and availability of cultured isolates. Moreover, to what extent such defined consortia can be successfully applied for *in situ* bioremediation remains uncertain (28, 29). In comparison, a top-down approach that relies on establishing a robust consortium through enrichment appears to have less of these concerning issues (30, 31).

Acetylene has wide inhibitory effects on microbial processes, i.e. methanogenesis, nitrogen fixation and reductive dehalogenation (32, 33). Recent studies revealed that acetylene hydratase and nitrogenase of *Pelobacter* sp. strain SFB93 can ferment and reduce acetylene, respectively, (34), abating the inhibition of acetylene on reductive dehalogenation of PCE to ethene when incubated with *Dehalococcoides* (35). However, it has not been shown to what extent this inhibitory effect of acetylene acts at transcriptional, translational or enzyme level. In addition, whether the coexisting nitrogenase in OHRB can prevent the inhibition of reductive dehalogenation still remains elusive.

In this study, we isolated a mutualistic consortium that is capable of reductive debromination of 2,6-DBP without adding external cobalamin, vitamin B12 (B12). Further genome-resolved analyses identified five novel bins, of which three were bearing RDase and TPh-RDase genes possibly involved in reductive debromination that was post-transcriptionally inhibited by acetylene. Metabolic pathway analysis confirmed that this consortium contains the complete *de novo* B12 biosynthesis pathway. Interestingly, further phylogenomic analyses suggested *Marinifilum* and *Ancylomarina* as new potential OHRB.

## Materials and methods

### Chemicals

PCE, 2,6-DBP, 2,4-DBP, 2,4-6-DBP, 1,4-dibromobenzene (1,4-DBB), 1,2-DBB, 1,3-DBB, 1,2,4-tribromobenzene (1,2,4-TBB), 2,6-dichlorophenol (2,6-DCP), 2,4-DCP, 2,4,6-TCP, 1,4-dichlorobenzene (1,4-DCB), 1,2-DCB, 1,3-DCB, 1,2,4-TCB, benzene, 2,4,6-triiodiphenol (2,4,6-TIP), 2,4-DIP, 2,6-DIP, 2-IP, and 4-IP were purchased from Sigma-Aldrich. Stock solutions of sulfate (0.5 M), sulfite (0.5 M), thiosulfate (0.5 M), nitrate (0.5 M), formate (0.5 M), acetate (0.25 M), pyruvate (0.5 M), and lactate (0.5 M) were prepared by filter sterilization (syringe filter, 0.2 µm, mdimembrane, Ambala Cantt, India). All other (in)organic chemicals were of analytical grade.

### Isolation and cultivation

Marine medium was prepared for cultivation as previously described (36, 37). The final growth medium was composed of 50 mL anoxic marine medium, Na_2_S·9H_2_O (0.48 g/L, 2 mM) serving as the reducing agent, and Resazurin (0.005 g/L) as the redox indicator. The headspace of culture bottles was exchanged with N_2_/CO_2_ (80 : 20%, 140KPa), and bottles were sealed with Teflon-coated butyl rubber septa and aluminum crimp caps (GRACE, MD, USA). Slant tubes contained 5 ml marine medium with 0.8% low-melting point agarose (Sigma-Aldrich) and were incubated in the dark at 20 °C. The isolation was initiated starting from the mother bottle, S.SD23 (third bottle of PCE dechlorination after two-time serial dilution with additional sulfate, details in Figure 2 of previous work (6)), and 200 µL was used to inoculate the anaerobic slant tube. Colonies were picked from the slant and transferred into liquid marine medium to check for PCE dechlorination in the presence of additional sulfate. The above-mentioned halogenated compounds were tested as the electron acceptors with lactate as the electron donor and carbon source. To study the growth of the community on 2,6-DBP (200 μM), hydrogen (20 mM) and acetate (5 mM) served as the electron donor and carbon source respectively. Acetylene (≈ 1.8 mM) was injected into the bottles and served as the inhibitor for microbial processes, including reductive dehalogenation. Prior to this growth measurement on H_2_/acetate as the sole electron donor and carbon source with 2,6-DBP as the electron acceptor, 2.5 ml actively debrominating culture was transferred three times (10% vol/vol) in a row to 48 ml fresh medium supplemented with H_2_/acetate and 2,6-DBP as described above and incubated to rule out the influence of residual medium components, especially lactate and sulfate. The generation of a B12 independent culture followed the same steps, i.e. three consecutive transfers and incubation to exclude residual B12 in the cultures.

**Figure 1.**
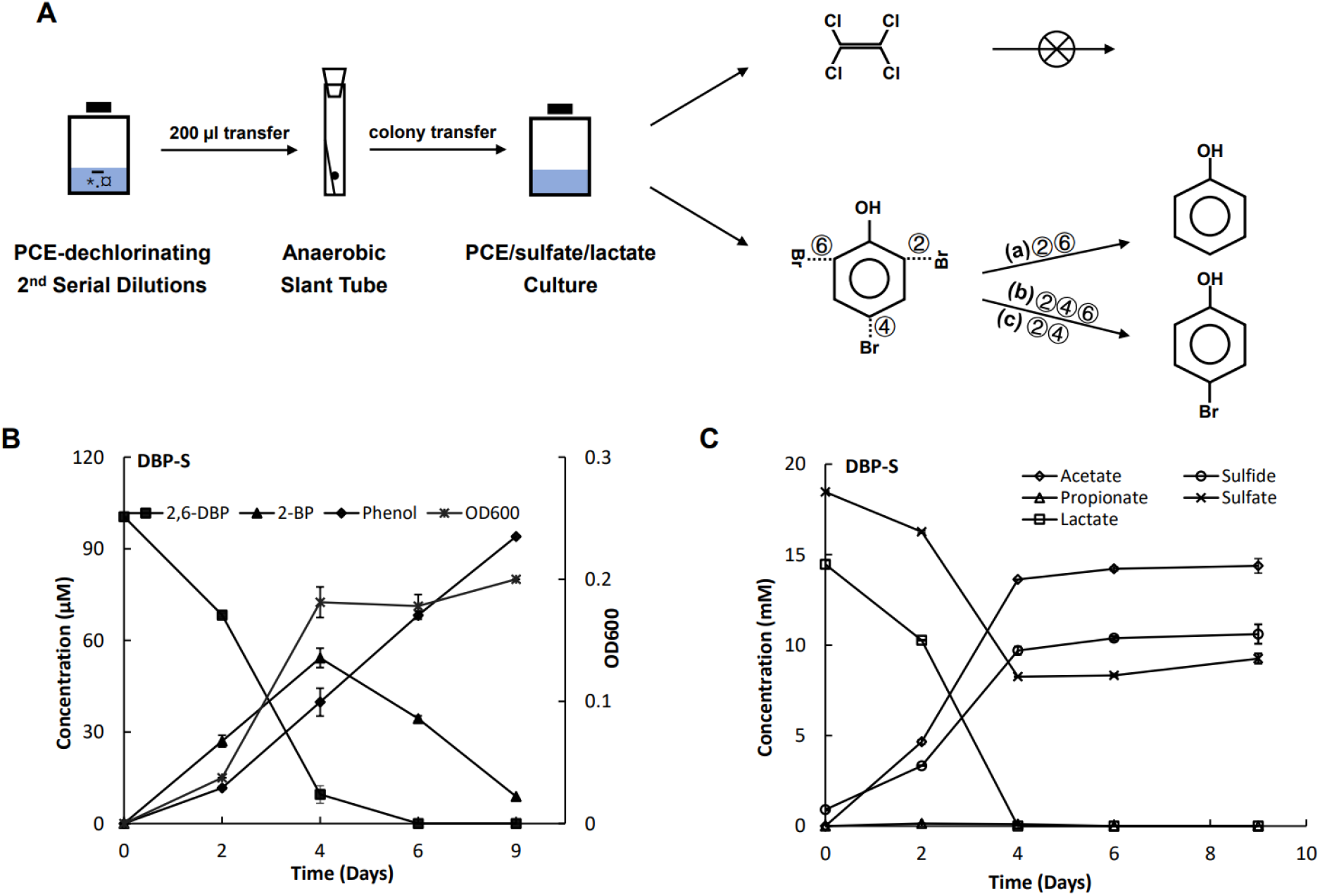
Schematic diagram of colony isolation and its reductive debromination of 2,6-dibromophenol (2,6-DBP) in the presence of sulfate. The mother bottle (10^-3^) was selected from the second serial dilution in the presence of sulfate as previously described (6). The colony-derived culture could completely debrominate 2,6-DBP to phenol, whereas 2,4,6- or 2,4-brominated phenols were transformed into 4-bromophenol (4-BP) (A). Debromination of 2,6-DBP to phenol (B) with the formation of 2-bromopenol (2-BP) as the intermediate, together with sulfate reduction to sulfide (C) to support the bacterial growth (OD600). Three replicate bottles were set for the reductive debromination of 2,6-DBP with additional sulfate. Data are presented as mean ± standard deviation (SD). Error bars indicate the SD.

**Figure 2.**
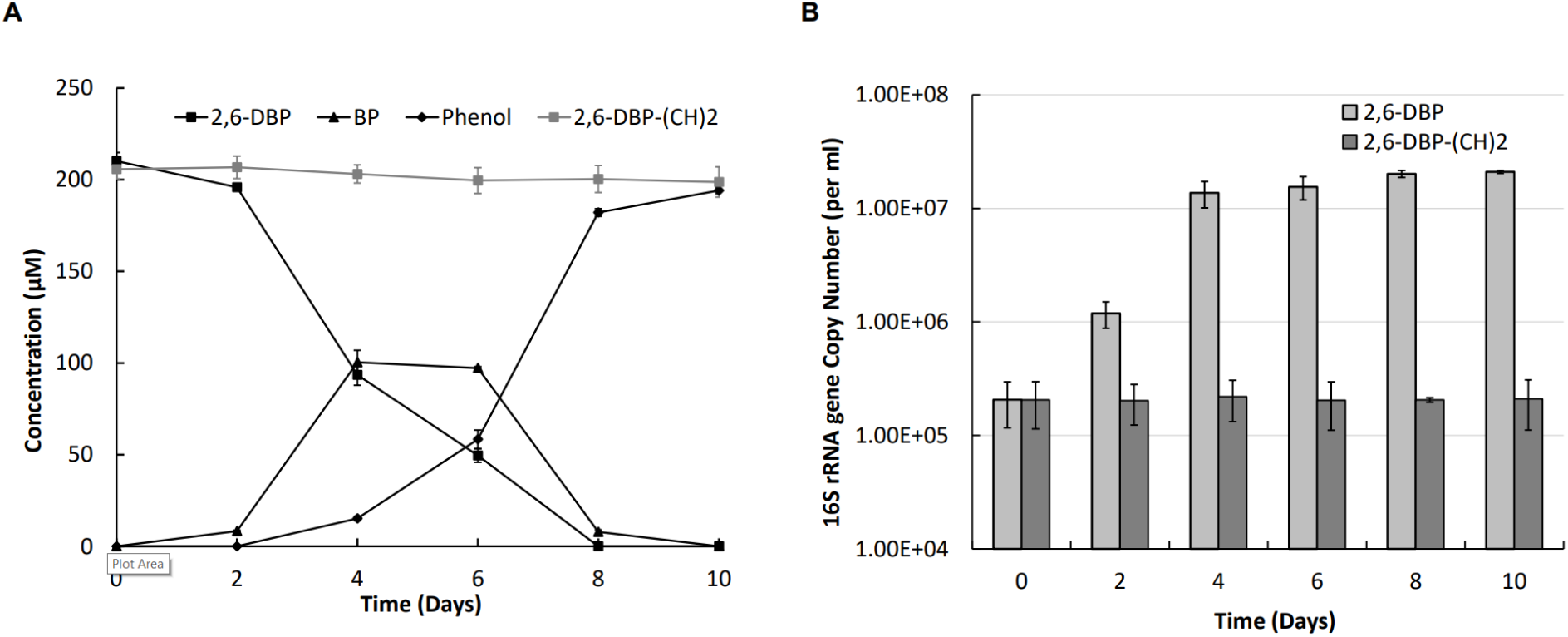
Reductive debromination to support the growth of consortium. H_2_/acetate served as the electron donor and carbon source for 2,6-DBP debromination, and acetylene added to the negative control (grey line) inhibited 2,6-DBP debromination (A). Quantitative PCR targeting the bacterial 16S rRNA gene was used to assess the cell growth with and without acetylene added (B). (CH)2: acetylene; three replicate bottles were set for the experiment, and data are presented as mean ± standard deviation (SD). Error bars indicate the SD.

### Analytical methods

PCE, trichloroethene (TCE), cis-dichloroethene (cDCE), trans-dichloroethene (tDCE), ethene and acetylene were detected by gas chromatography combined with mass spectrometry (GC-MS) with an Rt^®^-Q-BAND column (Retek, PA, USA) and DSQ-MS (Thermo Fisher Scientific). Hydrogen gas was detected by Compact GC (Global Analyzer Solutions, Breda, The Netherlands) with a pulsed discharge ionization detector (GC-PDD). Thermo Scientific Accela High-Performance Liquid Chromatography (HPLC) system installed with an Agilent Poroshell 120 EC-C18 column and a UV/Vis detector was used to measure halogenated phenols, and benzenes, benzene, and phenol. Organic acids were measured by SHIMADZU LC2030 PLUS coupled with a Shodex SUGAR Series® SH1821 column, including lactate, propionate, acetate and formate. Sulfate, sulfite, thiosulfate and nitrate were measured by using Thermo Scientific DionexTM ICS-2100 Ion Chromatography System (Dionex ICS-2100). Sulfide was analyzed by photometric method using methylene blue as described previously (Cline 1969).

### Scanning Electron Microscopy (SEM)

The culture was sampled for field emission scanning electron microscopy (FE-SEM). Five ml culture was sampled and incubated with glutaraldehyde (2.5%) for 20 min, and then prefixed to the cover slide coated with poly-L-lysine for 2 h. After that, the cover slide was washed three times with 0.1 M sodium cacodylate (pH 7.2) and then 1 h fixed with 1% osmium tetroxyde in the same cacodylate buffer. Finally, the samples were dehydrated with various ethanol concentrations and incubated for 10 min at each step. Imaging of the sample was completed using a Magellan 400 instrument at the Wageningen Electron Microscopy Center (WEMC).

### DNA and RNA extraction and reverse-transcriptase quantitative PCR (RT-qPCR)

The consortium cultures were first centrifuged at 10,000 × g for 5 min, the supernatant was discarded and the precipitates were then washed three times with 200 µL TE buffer (pH = 7.0, 4 °C) on ice to remove any residual medium components that might interfere with downstream DNA and RNA extraction. For genomic DNA extraction, MasterPureTM Gram positive DNA purification Kit (Epicentre, WI, USA) was used, following manufacturer’s instructions. For RNA isolation, prior to the sample collection, cultures in eighteen bottles were grown with additional lactate (15 mM) and sulfate (20 mM) to exponential phase for 72 h after transfer without any additional 2,6-DBP. Nine bottles were collected as the 0 hour samples. In the remaining nine bottles, three were kept as before without any injection, three replicate cultures were injected with 200 µM 2,6-DBP, and the last three were injected with 200 µM 2,6-DBP and with 1.8 mM acetylene together, and cells were collected and washed as described above. Collected biomass was mixed with 0.4 ml cold TE buffer (4 μl *2*-mercaptoethanol), and 0.5 ml TRIzolTM reagent (Thermo Fisher Scientific) was added, followed by bead-beating for 3 min (3 times, 1 min per time with cooling on ice in between) at speed 5.5 (FastPrep-24 5G, MP biomedicals, Irvine, CA, USA). After bead-beating, UltraPureTM phenol : chloroform : isoamyl alcohol in ratio of 25 : 24 : 1 in 200 μl (Thermo Fisher Scientific) was added and mixed by vortex. Then, the separated aqueous phase containing RNA was transferred to an RNeasy column (Qiagen, Venlo, The Netherlands) for purification followed by DNase I (Roche, Almere, The Netherlands) treatment to remove residual DNA. In order to check for purity of the obtained culture, a near-full length fragment of bacterial 16S ribosomal RNA (rRNA) gene was amplified using universal bacterial primers 27F/1492R (Table 1) and subjected to Sanger sequencing as previously described (Suzuki and Giovannoni 1996). Bacterial 16S rRNA gene-targeted qPCR was used for assessing the growth of the microbial community via the increase of 16S rRNA gene copy numbers using the general primers Eub341F/Eub534R (Table 1). RT-qPCR was introduced to measure the relative expression of RDase genes by using the One Step PrimeScript™ RT-PCR Kit (Perfect Real Time) (Takara Bio, Saint-Germain-en-Laye, Germany). Primers targeting RDase genes were designed using the NCBI primer design tool with the setting parameters, Expected PCR product sizes ranged from 75bp to 150bp and melting temperatures from 57 to 60 °C (Table 1). RT-qPCR targeting the bacterial 16S rRNA was used for normalization of RDase gene expression.

**Table 1.**
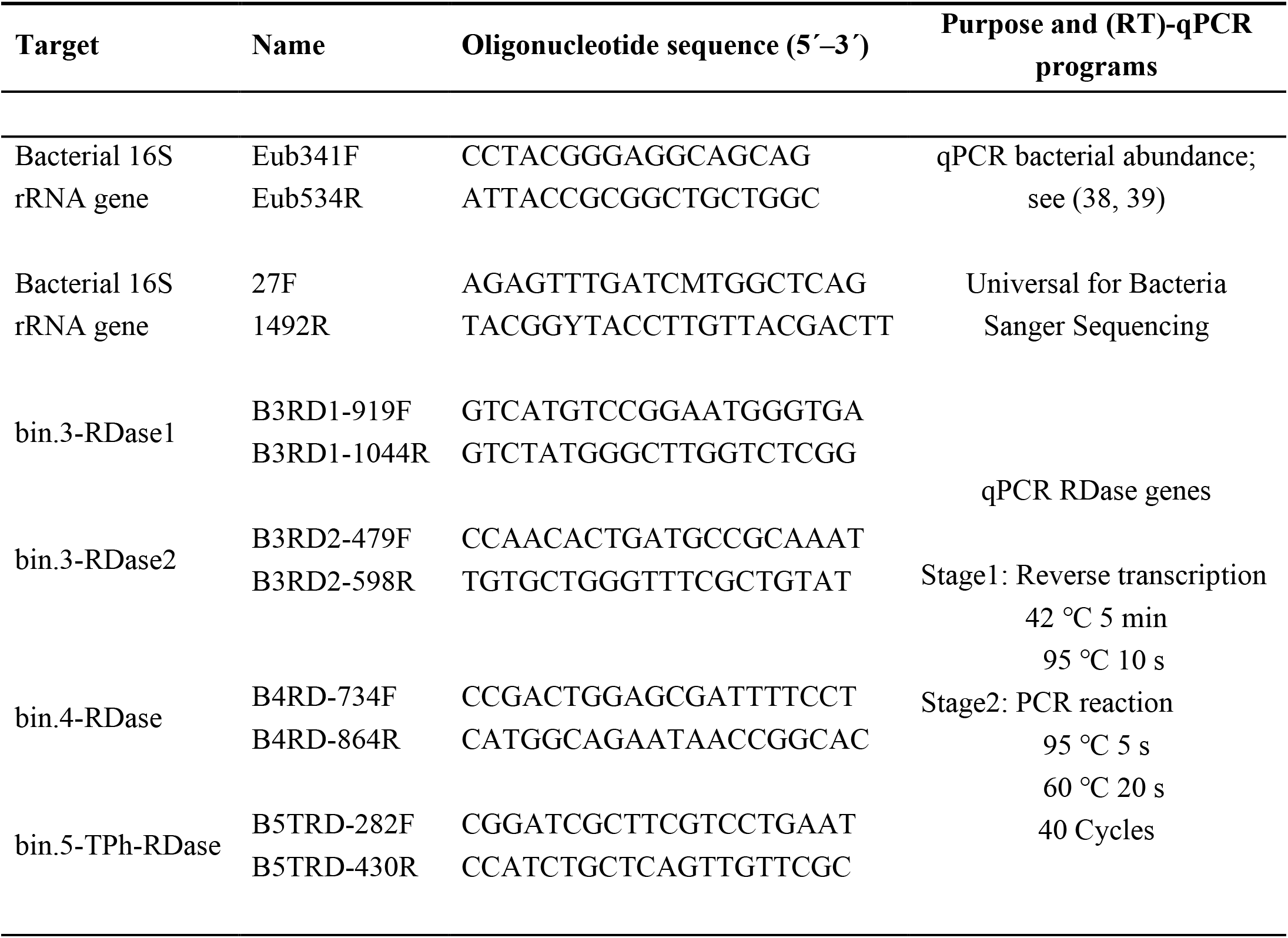
Primers used in this study for (RT)-quantitative PCR

### Metagenome sequencing and analyses

Metagenome sequence data was generated by Illumina paired-end short reads (PE150, Novaseq6000, Novogene) and PacBio long reads (Sequel, Novogene). The raw paired-end Illumina and PacBio reads were at first examined using the quality check module, “read_qc” of MetaWRAP (40). Low-quality reads as well as human contaminant reads were removed. The raw PacBio long-read data was combined to improve the quality of assembly following OPERA-MS (41). Considering the improved features of the hybrid assembly in comparison to the short-read assembly, including contigs’ N50, maximum contig size and number of contigs, the follow-up analyses were based on the hybrid assembly. “Kraken2” was used to identify the taxonomic composition of the community by mapping the clean Illumina reads against the SILVA database version 138 (42, 43) and to calculate the relative abundance of 16S rRNA gene reads of each taxonomy over the total reads of the 16S rRNA gene. The resulting hybrid assembly was binned and refined using the MetaWRAP modules, “Binning module” combining metabat2, maxbin2 and concoct with the cut-off set at > 50 % completeness and < 10 % contamination, and “Bin_refinement” to improve the bin set. The abundance of the refined bins was quantified using the “quant_bins” module, and the relative abundance of each bin was calculated by mapping the Illumina data to a concatenated file of the five bins using default parameters. Bins were reassembled, taxonomically classified and annotated via the “classify_wf” workflow of GTDB-Tk and the “prokka” module of MetaWRAP, respectively (44). The relative abundance of bins was calculated using CoverM v0.6.1 (https://github.com/wwood/CoverM).

### Phylogenetic analysis of bins and inference of metabolic pathways

There were five bins assembled, and their classification by GTDB-Tk revealed that all of them were new species as their average nucleotide identity (ANI) with entries in GTDB (v207) was lower than 95%. Fifty-two representative genomes most closely related to bins were extracted from the GTDB database (v207) and were included for phylogenetic analysis by GToTree (45). Metabolic traits associated with reductive dehalogenation were also inferred, including the metabolic genes encoding RDase (*rdh*), TPh-RDase (TPh-*rdh*), haloacid dehalogenase (*hdh*), pathways involved in the metabolism of sulfate, nitrate, hydrogen, nitrogen, acetylene, Wood-Ljundahl pathway (WLP), *de novo* B12 biosynthesis and B12 transporters (34, 46–54). To this end, MagicLamp (28) was used with the retrieved COG numbers of metabolic genes from the .gff format of bins and their related representative genomes to infer the metabolic potential of the consortium. The constructed phylogenetic tree was visualized and modified by ggtree (55).

## Results

### Debrominating potential of the isolated consortium

In order to isolate members of the stable, sediment-free PCE dechlorinating enrichment derived from Aarhus Bay sediment (6), we inoculated a sample of this culture onto an anaerobic agar slant (Figure 1A). We were able to retrieve a single colony that developed after 69 days of incubation. After transfer to liquid medium, the derived culture exhibited debrominating activity of 2,4,6-DBP and 2,6-TBP to 4-BP and phenol, respectively, but was no longer able to dechlorinate PCE (Figure 1A). To better understand the physiological traits of the colony-derived culture, different electron donors and carbon sources, such as H_2_/acetate and pyruvate, and various electron acceptors, such as sulfate and nitrate, and in particular a range of different halogenated compounds, were tested (Table 2). In line with the initial results, the culture only displayed debrominating potential rather than dechlorination or deiodination. While the colony-derived culture was able to utilize sulfate, sulfite, and thiosulfate as electron acceptor, it was not capable of reducing nitrate. The culture was found to completely debrominate 2,6-DBP to phenol after nine days with the formation of bromophenol as the intermediate (Figure 1B). Meanwhile, the added sulfate served as a competitive electron acceptor and was reduced to sulfide, in which lactate as the electron donor and carbon source was utilized with the production of acetate (Figure 1C). In line with what was observed for the incubation in the presence of sulfate, the reductive debromination also proceeded in the sulfate-free culture (Figure S1A), in which lactate was consumed fermentatively with the formation of propionate and acetate in a ratio of roughly 2 : 1 (Figure S1B).

**Table 2.**
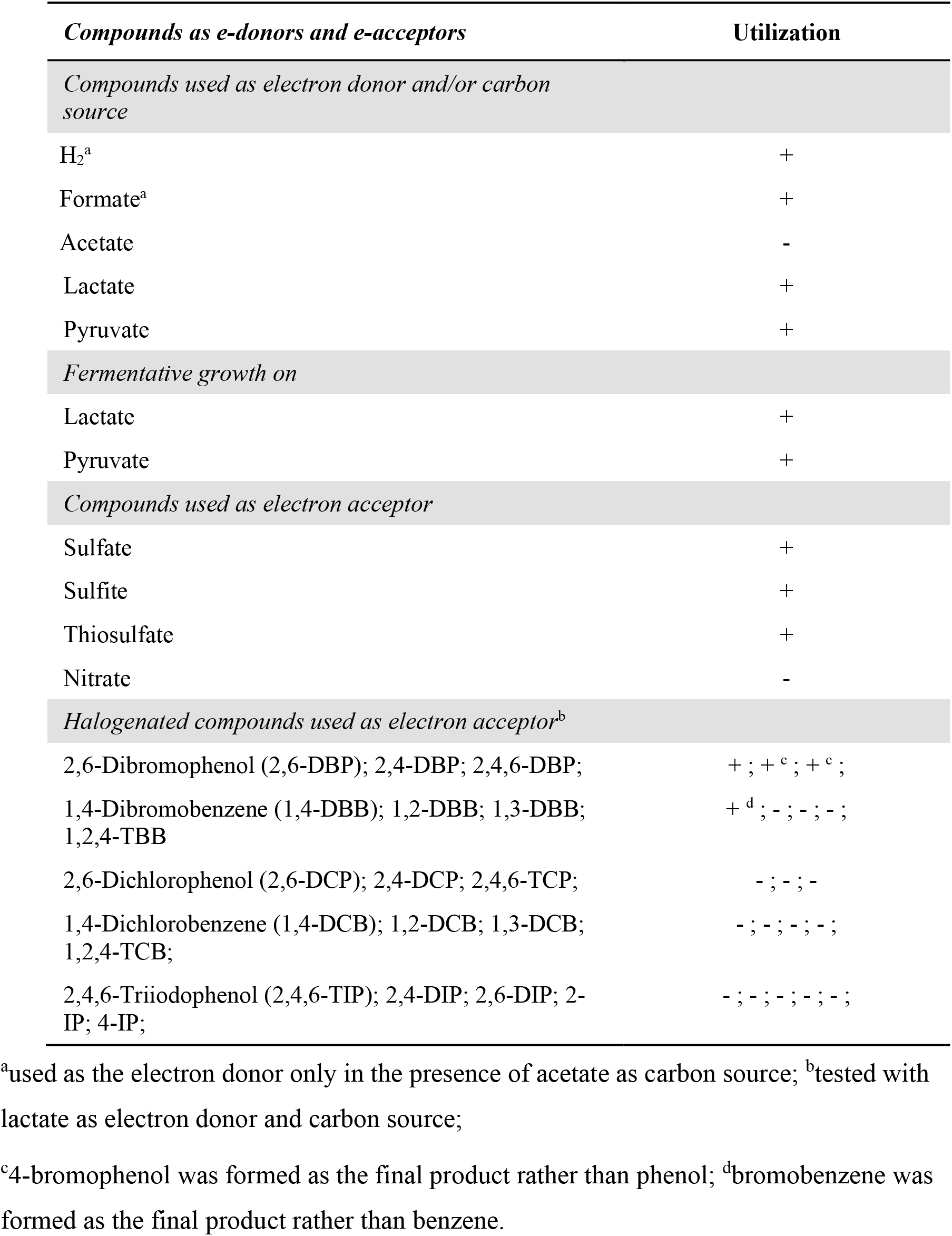
Physiological properties of the isolated consortium

In order to check for purity of the colony-derived culture, Sanger sequencing of a PCR product obtained using universal bacterial 16S rRNA gene-targeted primers 27F/1492R was attempted, however, the resulting sequence could not be resolved, indicating that the culture comprises a mixed consortium rather than a pure culture. Further scanning electron microscopy (SEM) analyses confirmed the presence of diverse morphologies in the community (Figure S1C), albeit with a rod-shaped cell-type being predominant.

### Bacterial growth supported by reductive debromination of 2,6-DBP

Reductive dehalogenation has previously been shown to be either catalyzed by respiratory corrinoid-dependent RDase or non-respiratory glutathione-dependent TPh-RDase (9, 11, 12, 56). To this end, reductive dehalogenation enabled by respiratory RDase as the terminal electron accepting process has been characterized to contribute to respiratory energy conservation for bacterial growth (56, 57). Previously, acetylene has been shown to inhibit reductive dehalogenation by *Dehalococcoides* (58). Our experiments validated acetylene can specifically inhibit RDase activity without influencing other metabolic activities, including lactate consumption and sulfate reduction (Figure S2). We therefore measured the bacterial growth during 2,6-DBP debromination compared to a culture to which acetylene was added. H_2_/acetate was added as the electron donor and carbon source and 2,6-DBP as electron acceptor to the cultures, respectively, in which the bacteria grew from 2.06 (± 0.90) × 10^5^ to 1.59 (± 0.16) × 10^7^ 16S rRNA gene copies / ml after the complete debromination of 200 µM 2,6-DBP in ten days (Figure 2). Simultaneously, in cultures to which acetylene was added reductive debromination of 2,6-DBP was completely stopped (Figure 2A), and the growth of the consortium was halted (Figure 2B). This finding suggested that one or more respiratory RDases were active as reductive debrominase of 2,6-DBP, contributing to the cell growth of the consortium. Intriguingly, cell growth came to a halt after four days with half of the added 2,6-DBP debrominated, suggesting that the non-respiratory RDase, TPh-RDase, might have also contributed to the degradation of 2,6-DBP without bacterial growth as the output. Indeed, additional research is needed to further investigate this observation.

### *Desulforhopalus*-dominated consortium for reductive debromination

In order to further investigate the composition and metabolic potential of the consortium, metagenome sequencing, including short-read and long-read sequencing, was employed. Taxonomic assignment of 16S rRNA gene sequences derived from the metagenome dataset revealed that *Desulforhopalus*, classified into *Desulfobacterota*, accounted for the largest percentage with 29%, followed by populations of *Klebsiella* and *Myroides* with 8% each, and genera *Mycoplasma* (7%) and *Endomicrobium* (7%) (Figure 3A). In addition, 11% of the sequences were affiliated with the *Bacterodia,* while 10% belonged to *Desulfobacterota* other than *Desulforhopalus*. This overall aligned well with the community analyses of the original PCE-dechlorinating cultures (6). To accurately decipher the taxonomy and genomic information, short and long metagenome sequence reads were assembled and binned into five genomes (MAGs) that were further classified using the GTDB-TK workflow. bin.1 was found to belong to the genus *Oceanispirochaeta*, bin.2 and bin.3 were affiliated to *Desulfoplanes*, bin.4 to the family *Marinifilaceae*, and bin.5 was affiliated with the genus *Desulforhopalus* (Table S1). All five MAGs were identified as new species based on ANI values with most closely related species being lower than 95%. The reads of the five MAGs accounted for 95.96% of total reads, with bin.5 accounting for 65.62%, bin.3 13.61%, bin.2 10.56% and bin.4 for 3.55%, leaving 4.04% of all reads unmapped (Figure 3B). Furthermore, bin.5 was characterized by 100% completeness and 0% contamination according to 120 bacterial single-copy maker genes. In summary, both approaches supported the notion that bin.5, identified as member of *Desulforhorpalus*, was the most abundant member of the consortium.

**Figure 3.**
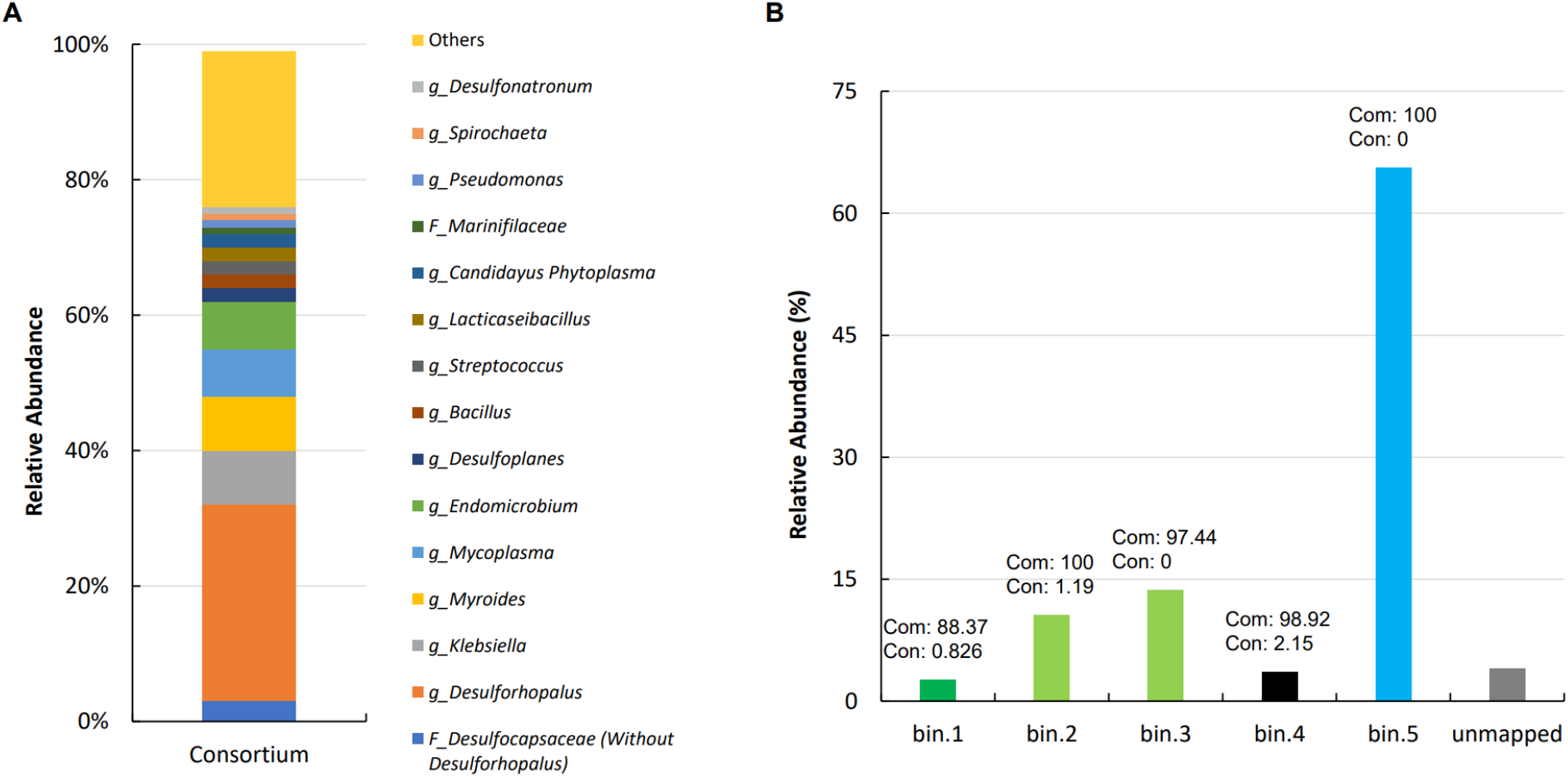
Relative abundance of microbial community members and the assembled bins within the colony-derived consortium. Microbial composition was calculated based on the mapping file generated from the clean raw Illumina reads against the SILVA database version 138 (42, 43)(A). The hybrid assembly was binned into five novel bins, and the relative abundance of each bin was calculated based on its numbers of reads over the numbers of total reads (B). “Others” represent the assigned taxa with a relative abundance < 1% and unassigned taxa; *F*, family; *g*, genus. Com: completeness (%); Con: contamination (%); unmapped: reads were not binned.

### RDase genes and their expression during reductive debromination

To further examine, which of the bins are responsible for the observed reductive debromination, MAGs were annotated, leading to the identification of three genes predicted to encode canonical corrinoid-dependent RDases, two of which were found on bin.3 and one on bin.4. A fourth gene, predicted to code for a thiolytic glutathione-dependent RDase was found on bin.5 (Figure 4A). Subsequently, the genomic context, i.e. the two neighboring genes upstream and downstream of each RDase gene, was included in the analysis. All three RDase genes encoding the catalytic subunit of respiratory RDases (RdhA) were accompanied by genes predicted to encode the cognate membrane anchor protein (RdhB) Interestingly, both RDase genes found on bin.3 were accompanied by a gene predicted to code for a formate hydrogenase transcriptional activator, HyfR (59), suggesting both loci originated from the same ancestor. Another transcriptional regulator, Btr (60), for siderophore bacillibactin production under iron-limiting conditions, was found encoded upstream of RDase in bin.4. Finally, upstream of the TPh-RDase encoding gene we observed a gene predicted to code for a transcriptional regulator, NoDD2, regulating nodulation factor production (61). Although these different regulators have been described with divergent functions, their roles are in general closely related with energy conservation and nutrient acquisition. As to the three predicted corrinoid-dependent RDases, three commonly conserved functional motifs were found, including a twin arginine signal peptide (RR), as well as two iron-sulfur cluster motifs, FCXXCXXCXXXCP and CXXCXXXCP, that were conserved in sequence and structural configuration for binding the iron-sulfur clusters to transfer electrons (Figure 4B). TPh-RDase was classified into the superfamily of glutathione-S-transferases (GST) that differ from respiratory RDase in sequence and conserved motifs. Sequence alignment of TPh-RDase in bin.5 to that of *Sphingobium chlorophenolicum* revealed that two catalytic site residues, cysteine (C) and serine (S), were conserved in sequence and simulated structure. All four RDases were predicted to be membrane-spanning enzymes, suggesting direct contact with substrates and dehalogenation in the extracytoplasmic space (Figure S3).

**Figure 4.**
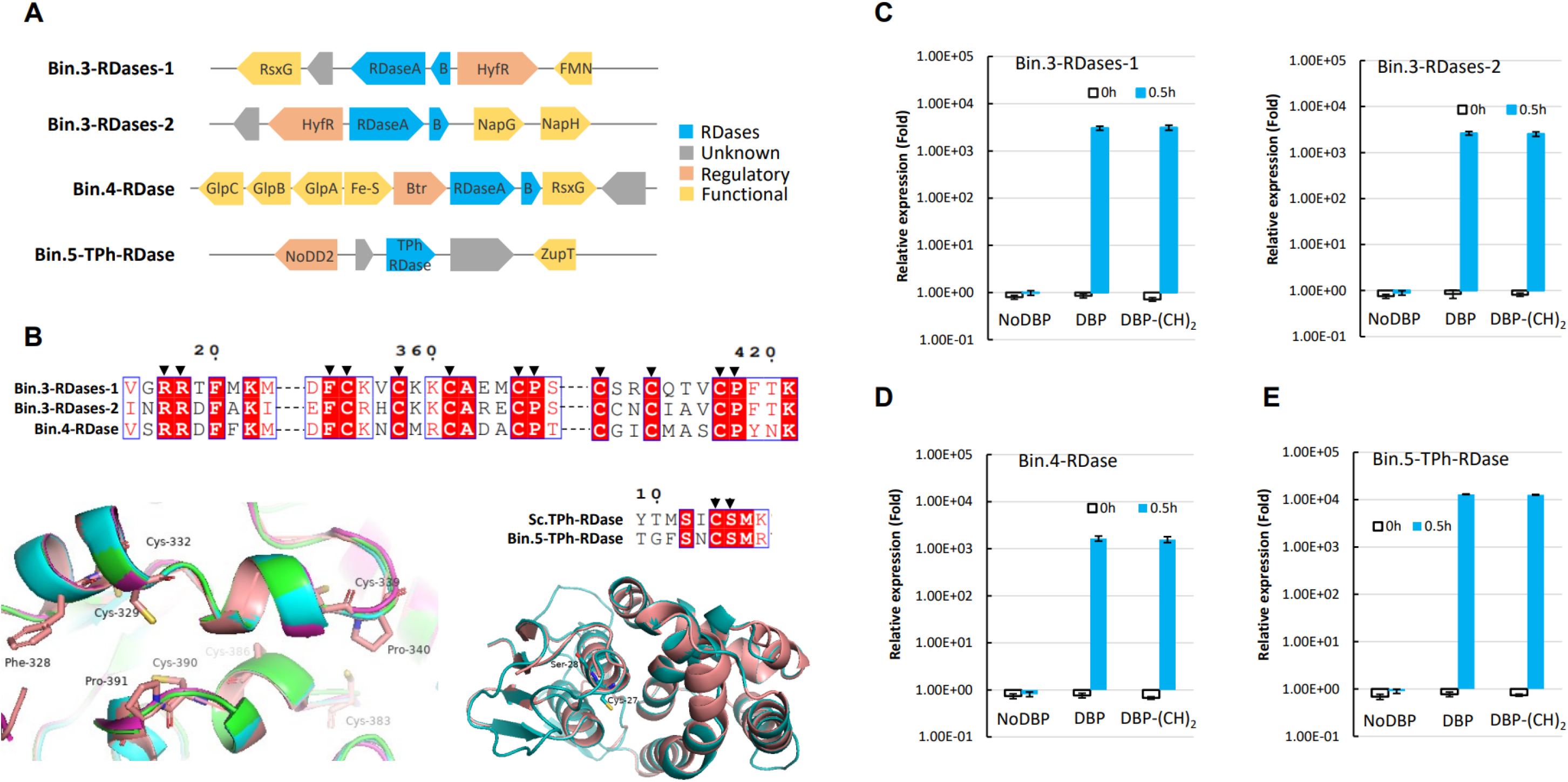
The genomic context, conserved motifs and structures, and expression of RDase encoding genes identified in the bins. RDase genes with the two neighboring genes located up- and downstream (A). Amino acid sequence alignment of conserved motifs of the respiratory RDases of bin.3 and bin.4, and TPh-RDase of bin.5 and *Sphingobium chlorophenolicum* respectively displayed by EsPript 3.0, and corresponding superimposed structures as generated by PyMol (v.2.3.4) (see below for additional details) (B). Reverse-transcriptase quantitative PCR was used to measure the expression of RDase genes from bin.3 (C) and bin.4 (D), and TPh-RDase gene from bin.5 (E), before (0 h) and 30 min (0.5 h) after addition of 2,6-DBP only, no addition, and addition of 2,6-DBP and acetylene together. Expression values were normalized to 16S rRNA measurements. For the structures shown in B, respiratory RDase (PDB: 5m2g1, in salmon) from *Sulfurospirillum multivorans* was aligned as the best structural template (18). In the superimposed structure, bin.3-RDase1 is depicted in green, bin.3-RDase2 in cyan and bin.4-RDase in magenta. TPh-RDase (PDB: 7za5.1, in salmon) is shown as dimer as the structural basis for TPh-RDase of bin.5 in teal. FMN: Flavoprotein that can bind the flavin mononucleotide (FMN) as the prosthetic group; RDases: composed of RDaseA and anchor protein B; HyfR: transcriptional regulator of formate hydrogenase; RsxG: electron transporter; NapGH: electron transporter associated with periplasmic nitrate reductase. GlpABC: Anaerobic glycerol-3-phosphate dehydrogenase complex; Btr: transcriptional regulator; Fe-S: Ferredoxin with iron-sulfur clusters; TPh-RDase: none respiratory RDase; NoDD2: regulator for nodulation; ZupT: zinc transporter. Arrows indicate the conserved sites, in which some were labelled in the superimposed the structure. NoDBP: no 2,6-DBP added; DBP-(CH)2: both 2,6-DBP and acetylene added. Each treatment in C-E was set with three replicates and values represent mean ± standard deviation (SD). Error bars indicate the SD.

As a next step, we analyzed the transcription of RDase genes using RT-qPCR. This analysis revealed that expression of all four RDase genes were induced several orders of magnitude within 30 min after the addition of 2,6-DBP. More specifically, the relative expression of the TPh-RDase gene increased up to 1.30 (± 0.16) × 10^4^ fold compared to the control to which no organohalide was added. Similarly, expression of genes encoding the respiratory RDases, i.e. RDase-1 and RDase-2 of bin.3, and RDase of bin.4, increased by 3.04 (± 0.29) × 10^3^, 2.63 (± 0.24) × 10^3^, and 1.65 (± 0.20) × 10^3^ fold, respectively (Figure 4C). Interestingly, we found that the inhibitor acetylene did not affect RDase gene expression, suggesting that inhibition acts post-transcriptionally.

### Phylogenomic analyses of bins and closely-related representative genomes

To gain a better understanding of the phylogenies and metabolic traits of bins compared to genomes of closely-related organisms, we constructed phylogenetic tree of bins and associated representative genomes, and subsequently investigated encoded metabolic traits related to the above-mentioned physiological data (Figure 5). As mentioned above, these MAGs were classified into four taxonomical groups, in which bin.1 was affiliated with the genus *Oceanispirochaeta*, bin.2 and bin.3 were assigned to the genus *Desulfoplanes*, bin.4 was affiliated to the family *Marinifilaceae*, and bin.5 was assigned to the genus *Desulforhopalus*. The 15 representative genomes from *Desulforhopalus*, including bin.5, all bear the conserved metabolic genes for sulfate reduction, *de-novo* corrinoid biosynthesis, H_2_ metabolism, and Wood-Ljungdahl pathway (WLP). Interestingly, whereas bin.5 was predicted to encode a glutathione-dependent TPh-RDase, the genomes of three other *Desulforhopalus* members contained genes predicted to code for corrinoid-dependent respiratory RDases, suggesting that these strains might have OHR potential. Furthermore, whereas most *Desulforhopalus* spp. genomes included in our analysis encoded the complete WLP, two members, *D. vacuolatus* and bin.5 were found to lack the core genes of the WLP, including *acsA*, *acsB* and *acsCD*, coding for carbon monoxide dehydrogenase, acetyl-CoA synthase and corrinoid iron sulfur protein (CFeSP), respectively (62). Interestingly, the genome of one of the isolates, *D. singaporensis,* encodes all genes for the complete WLP indicating the co-occurrence of OHR potential with the assimilation of C1 compounds. In contrast, bin.1, bin.2, bin.3 and bin.4 and their associated reference genomes were all found to lack the core genes for WLP, suggesting their incapability of one-carbon compound fixation.

**Figure 5.**
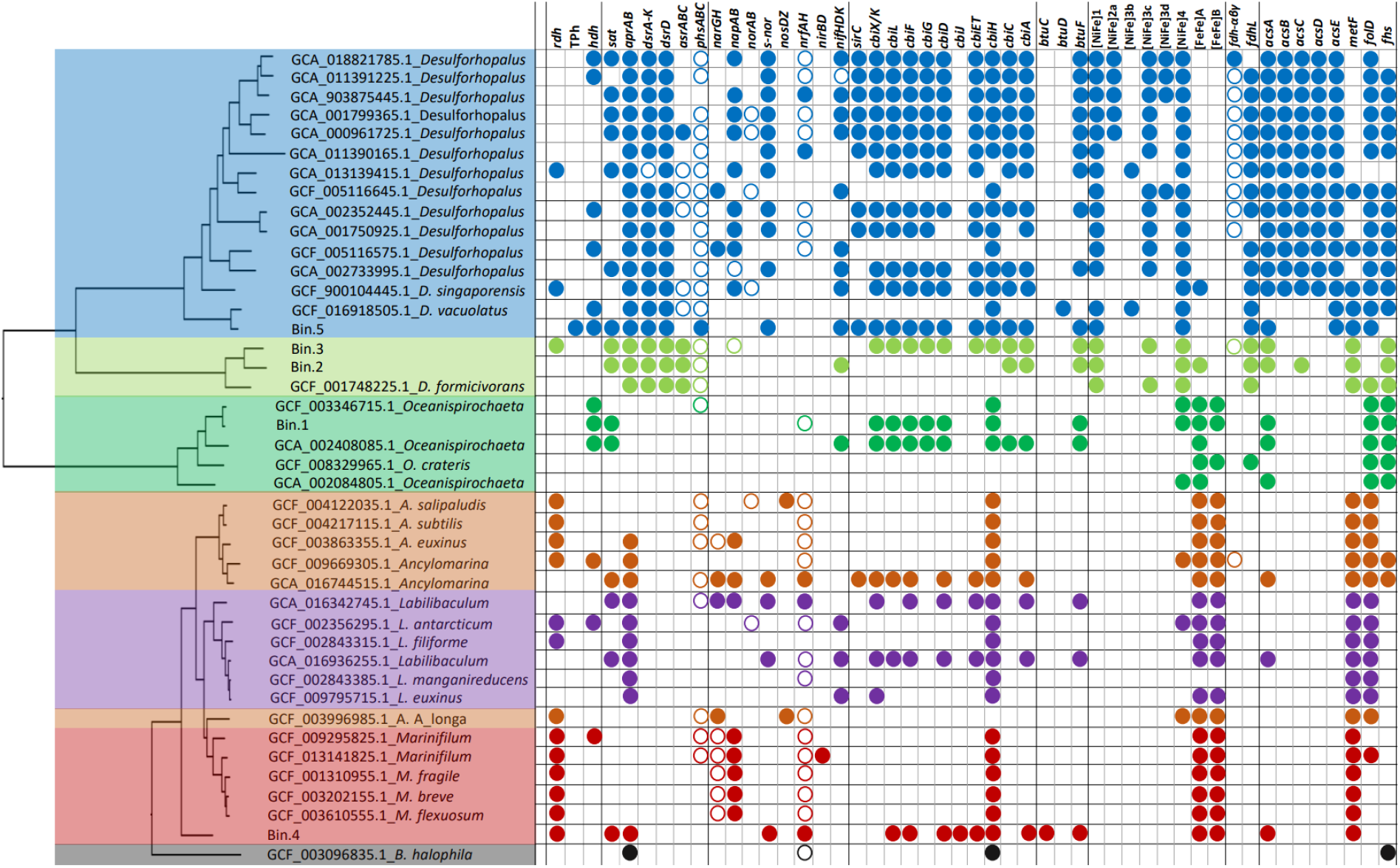
Phylogenies of the assembled bins and most closely related reference genomes, and comparison of their metabolic genes. In addition to the five bins, 52 related representative genomes from the GTDB database (v207) were included for tree construction by GToTree following the default parameters (45). Filled circles represent encoded functional genes or complete gene clusters. Open circles represent genes for which not the entire cluster was detected. Colors indicate the different phylogenies of bins and their closest relatives. *D. formicivorans*: *Desulfoplanes formicivorans*; *rdh*: RDase gene; TPh: TPh-RDase gene; *hdh*: haloacid dehalogenase gene; *sat*, *aprAB*, *dsrA-K*(*ABCTMK*), *dsrD*, *asrABC*, and *phsABC* for sulfate reduction (54); *narGH* and *napAB* for nitrate reduction (46); *norAB*, *nirBD*, and *nosDZ* for denitrification (64); *nrfAH* for ammonification (63); from *sirC* to *cbiA* for the complete anaerobic *de novo* corrinoid biosynthesis (51). *btuFCD* encodes the ABC-type corrinoid transporter (65); Four groups of [Ni-Fe] hydrogenases, 1, 2a, 3b, 3c, 3d, and 4, and two groups of [Fe-Fe] hydrogenases, A and B (48, 66); *fdh-αβγ*, or *fdhABG*, codes for formate dehydrogenase (50, 67); *fdhL* codes for NAD^+^-dependent formate dehydrogenase (68, 69); *acsABCDE*, *folD*, *metF* and *fhs* are critical genes for the Wood-Ljungdahl pathway (WLP) (70, 71).

B12 is the required cofactor for corrinoid-dependent RDases linked with OHR (25). Based on the analysis of bins, the *de novo* biosynthesis observed for the colony-derived consortium could be achieved by bin.3 and bin.5 separately, without the non-essential *cbiJ* gene, in line with what we observed earlier in marine *Desulfoluna* strains capable of OHR (37), or coordinatively achieved with bin.4 by providing the *cbiJ* gene missing in other bins. Hence, in summary, the genomic inferences in terms of predicted metabolic capacity of the colony-derived consortium sufficiently explain the observed OHR capacity of the consortium in the absence of added B12 (Figure S4).

All six genomes affiliated with the genus *Marinifilum*, including bin.4, were predicted to encode respiratory corrinoid-dependent RDases. Similarly, five out of six genomes affiliated with the closely related genus *Ancylomarina* were predicted to code for corrinoid-dependent RDases. Accordingly, these genera, as a subgroup within the *Marinifilacease* family, are predicted to comprise novel OHRB taxa. In addition, all included genomes of these taxa contained genes encoding type A and type B [Fe-Fe] hydrogenases for catalyzing H_2_ metabolism, in contrast to *Desulforhopalus* spp. genomes that were predicted to encode [Ni-Fe] hydrogenases. Interestingly, as outlined above, the critical *cbiH* gene of *de novo* B12 biosynthesis (51), was found conserved among the members of both genera *Ancylomarina* and *Marinifilum* suggesting a symbiotic role in *de novo* B12 biosynthesis. Being closely related to *Ancylomarina* and *Marinifilum*, available genomes of the genus *Labilibaculum* were found to contain similar functional genes, such as genes coding for [Fe-Fe] hydrogenases and CbiH, but only two of the six genomes included here, namely *L. antarcticum* and *L. filiforme*, were predicted to encode corrinoid-dependent RDases.

Physiological characterization of the consortium showed that the community was unable to reduce nitrate as the terminal electron acceptor, most probably due to the lack of genes encoding membrane-bound and periplasmic nitrate reductase (NarAB and NapAB) in these assembled bins. Similarly, the five bins retrieved from the consortium also were found to lack *norAB*, *nosDZ*, and *nrfAH* genes, indicating their inability of denitrification and ammonification, respectively (63, 64). Intriguingly, most of the reference genomes from the genera *Desulforhopalus* and *Marinifilum* were found to contain *napAB*, suggesting their potential to reduce nitrate to nitrite. Further, genes encoding the complete nitrogenase complex for nitrogen fixation were found in bin.2 and bin.5, suggesting their capability of fixing molecular nitrogen to produce ammonia in case of N-shortage.

## Discussion

Research described in previous work supported the notion that microorganisms inhabiting marine sediments of Aarhus Bay were capable of using a variety of halogenated compounds, such as PCE and 2,6-DBP, as terminal electron acceptors for anaerobic respiration (6). Henceforth, we set out to isolate the microorganisms responsible for this activity. This led to the isolation of a highly enriched consortium from an anoxic slant tube culture, which was characterized to dehalogenate brominated compounds rather than the chlorinated and iodinated compounds that the original enrichment was able to dehalogenate. The combination of short- and long-read metagenome sequence data generated in this study revealed that a population most closely affiliated with the genus *Desulforhopalus* was most abundant in this consortium, and five novel high quality MAGs could be assembled with > 85% completeness and < 2.5% contamination. Among these five MAGs, bin.3 and bin.4 were affiliated with the genus *Desulforplanes* and family *Marinifilaceae* respectively, and were identified as potential OHRB based on the presence of respiratory RDase-encoding genes, whereas, the *Desulforhopalus* associated bin.5 was found to encode a non-respiratory TPh-RDase. Intriguingly, the transcription of all RDase genes was strongly induced when 2,6-DBP was added, indicating that multiple strains simultaneously acted on this substrate, rather than outcompeting each other, at least for the duration of our studies that included multiple transfers. In addition, the consortium was found to have the genetic capacity for complete *de novo* B12 biosynthesis suggesting a symbiotic relationship for self-supplementing B12 to OHR.

This consortium was isolated from a stable, sediment-free PCE dechlorinating enrichment culture propagated in the presence sulfate, and which was mainly composed of members of the genera *Desulfoplanes*, *Desulfobacter*, *Bacillus* and *Desulforhopalus* (6). Of interest, the consortium lost the ability to dechlorinate PCE, but instead retained the capability to debrominate 2,6-DBP concomitant with retaining populations of *Desulfoplanes* and *Desulforhopalus* in the consortium, suggesting their role as potential debrominators, rather than being involved in the dechlorination observed in the original culture. This was further supported by the fact that, based on the metagenomic analyses of the consortium, both populations comprised a large proportion within the 2,6-DBP debrominating consortium described in this study (Figure 3), in which bin.3 classified into *Desulfoplanes* encoded two RDase genes, whereas bin.5 belonging to *Desulforhopalus* coded for a TPh-RDase, which was formerly characterized to catalyze glutathione-dependent thiolytic reductive dehalogenation in strictly aerobic bacteria (11). In contrast, our data suggest that the thiolytic RDase in bin.5 might function in anaerobes, and thus in anoxic environments. The sequence alignment and structural simulation of RDases reinforced their membrane-bound property (Figure S3), and conservation of catalytical motifs or residues (Figure 4). Interestingly, cysteine residues are essential and conserved in both types of enzymes. In corrinoid-dependent RDases a cysteine residue is recruited to bind B12, whereas in TPh-RDases the cysteine residue has been shown to form a covalent intermediate with glutathione during catalysis (72, 73). Acetylene was previously reported as a general inhibitor for many biological processes, such as methane production and fermentation, and OHR (58, 74). Therefore, we decided to assess whether we can use it in this study for the measurement of bacterial growth from OHR. To this end, our physiological data revealed that acetylene specifically inhibited reductive dehalogenation without inhibitory effects on sulfate reduction and lactate oxidation (Figure S2). Interestingly, we found bin.2 and bin.5 encode complete nitrogenase complexes (N_2_ases), NifHDK, which can reduce acetylene into ethene (75). However, there was no ethene conversion from acetylene suggesting that N_2_ases were inactive, possibly due to the presence of ammonium in the marine medium (9 mM), which can completely inhibit N_2_ase activity (76). Interestingly, recent studies revealed acetylene hydratase can allow bacteria to utilize acetylene as carbon source to support OHR (34, 35). Nevertheless, we did not find this enzyme being encoded across the genomic assembly of the consortium. Notably, the inhibition by acetylene occurred at post-transcriptional level as the transcription of all RDase genes was highly-induced after adding 2,6-DBP (Figure 2). Unfortunately, the exact mechanism of acetylene inhibition still remains unresolved. It was reported that acetylene can target metalloenzymes, especially the metal cofactors Fe, Ni, Mo, and Cu, or bind directly with substrates and/or active sites (75). It is thus tempting to speculate that the acetylene could target the iron-sulfur clusters or the catalytic sites of RDases to block the transfer of reducing equivalents, impeding RDase catalysis. Nevertheless, further experiments will be needed to elaborate the mechanism.

Respiratory reductive dehalogenation catalyzed by corrinoid-dependent RDases has been shown to serve as the terminal electron accepting process to complete the electron transfer through a membrane-associated electron transport chain (ETC) coupled to proton motive force formation and ATP production (18, 77, 78). Based on the five MAGs identified in this study, we propose ETCs and related activities in the 2, 6-DBP debrominating consortium studied here (Figure 6). bin.3 harbors two RDases, RDase1 and RDase2 (Figure 4), and we assumed that both RDases are part of ETCs that allow usage of hydrogen or lactate as electron donor as observed for *Dehalobacter restrictus* and *Desulfoluna spongiiphila*, respectively (37, 79, 80). The membrane-bound flavoprotein, binding flavin mononucleotide (FMN) that was found encoded adjacent to RDase1, is predicted to function in a similar way as RdhC in the electron transfer (81), which was initially hypothesized as transcriptional regulator of OHR in *Desulfitobacterium dehalogenans* (82). In addition to the FMN encoding gene, the RDase1 encoding gene is also accompanied by a gene coding for RsxG, homologous to RnfG, which has been shown to transfer electrons from the quinol pool to nitrogenase (83). Interestingly, a quinol dehydrogenase complex, NapGH, was predicted to be encoded downstream of RDase2 (16, 84), acting as electron carrier to pass the electrons from the menaquinol pool to RDase2. The NapGH complex usually transfers electrons to the NapAB complex for nitrate respiration (84). Based on the MAGs identified in this study, the consortium lacks the NapAB complex, in line with the observation that the culture was not able to use nitrate as electron acceptor (Table 1). With lactate added as the organic carbon source and electron donor, the fact that OHR and sulfate reduction occurred simultaneously, indicated the electrons were shared (Figure 6A). Lactate was consecutively oxidized into pyruvate and acetate by lactate and pyruvate dehydrogenase, respectively, to release the electrons entering the menaquinol pool. It is tempting to speculate that part of the electrons from this pool are transferred to the sulfate reduction pathway. To this end it should be noted that we did not observe genes coding for QmoABC that was previously shown to enable the conversion of APS to sulfite (54), and thus an alternative complex is needed to fulfil this role. Dsr proteins subsequently reduce the sulfite to sulfide. In the absence of sulfate, hydrogen was observed as the result of lactate fermentation, which then served as the intermediate electron donor that is oxidized by Ni-Fe hydrogen uptake-type hydrogenases (HupLS) to transfer the electrons to RDases through the above-mentioned electron carrier proteins. The ETC including the RDase encoded on bin.4 is proposed to function in a similar manner as outlined for bin.3 to generate and transport electrons for the generation of a proton motive force and ATP synthesis (Figure 6B). Intriguingly, an anaerobic glycerol-3-phosphate (G3P) dehydrogenase complex (GPDH), GlpACB, which was previously characterized as the membrane-anchoring enzyme to catalyze G3P conversion into dihydroxyacetone phosphate achieving the electron flow through the cytosol to the membrane (85), was found encoded on bin.4. We therefore speculated that this GPDH complex transfers electrons to the RDase initiating the ETP for OHR in this organism. Based on the model proposed here, further studies will be needed for the characterization and experimental validation of ETCs in the organisms studied here. In addition to the active respiratory RDases, the gene encoding the non-respiratory TPh-RDase from bin.5 was also highly expressed, suggesting an active role of this enzyme in the observed debromination of the 2,6-DBP to phenol (Figure 4E). The complete inhibition of reductive debromination by acetylene furthermore suggests that acetylene inhibited activity of both types of RDases. Finally, the *Desulforhopalus* population (bin.5) is proposed to act as the B12 provider for OHRB (bin.3 and bin.4) due to its nearly-complete gene-set for *de novo* B12 biosynthesis except for *cbiJ* (Figure 6C), which was not required for OHR as previous described (37). Moreover, bin.4 contains a *cbiJ* gene indicating that this might play a role in the complete B12 biosynthesis of the isolated consortium.

**Figure 6.**
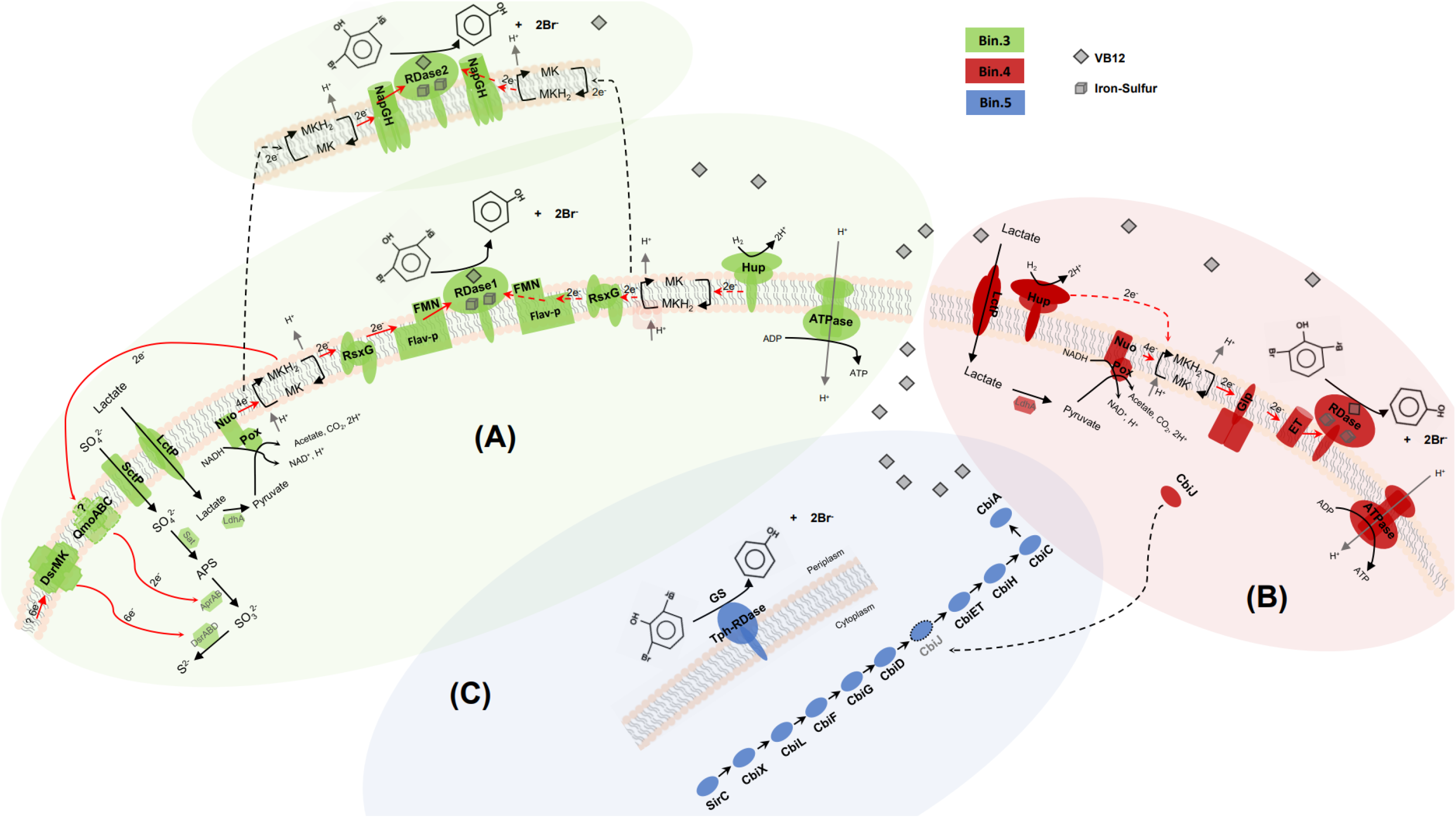
Proposed schematic overview of electron transport chains (ETCs) of organohalide respiration associated with sulfate reduction and de novo B12 biosynthesis. Electron transfer from lactate or H_2_ to the RDases and other proteins encoded on bin.3 to catalyze OHR and sulfate reduction (A). RDase of bin.4 follows a similar ETC pattern without sulfate reduction (B). In contrast, non-respiratory TPh-RDase of bin.5 transforms 2,6-DBP to phenol and provides the essential B12 for OHR to respiratory RDases (C). Color patterns used in this figure are in line with Figure 5. MKH_2_/MK: menaquinones; NapGH: periplasmic nitrate reductase; Flav-p: flavoprotein, which can bind with flavin mononucleotides (FMN); RsxG: electron transporter, homologous with RnfG (83); Nuo: NADH dehydrogenase; Pox: Pyruvate dehydrogenase; LctP: lactate permease; LdhA: lactate dehydrogenase; SctP: sulfate permease; Sat: sulfate adenylyltransferase; AprAB: APS reductase; DsrABD: dissimilatory sulfite reductase; DsrMK: electron transport complex function with DsrABD; QmoABC: electron transport complex; Hup: [Ni-Fe] hydrogen uptake-type hydrogenases, includes the large and small subunits; Glp: glycerol-3-phosphate (G3P) dehydrogenase complex; ET: ferredoxin as intermediate electron transporter; SirC-CbiA: complete B12 de novo biosynthesis. Red arrows indicate the assumed electron flow, black dotted arrows indicate parallel electron flow in (A), and CbiJ as the supplementary from bin.4 (B) to bin.5 (C) to form a complete B12 biosynthesis pathway; “? 6e^-^”: unknown source of deprived electrons; “? QmoABC”: not found in genomic annotation, and could be replaced by another functionally similar complex;

In summary, we provide evidence that the debromination potential of the isolated consortium is associated with the respiratory RDases and TPh-RDase encoded on bins that were phylogenetically-affiliated with potentially novel species within the genera *Desulfoplanes*, *Marinifilum* and *Desulforhopalus*. Moreover, phylogenomic analyses inferred for the first time that members of *Marinifilum* and *Ancylomarina* are potential OHRB, reinforcing a yet underestimated diversity of OHRB in marine environments. In addition, acetylene exerted specific inhibition on reductive dehalogenation post-transcriptionally, and could serve as an indicator to clarify the relationship between OHR, nitrogen fixation and acetylene oxidation in ammonium-low or free environments. Furthermore, the distribution of genes encoding a complete *de novo* B12 biosynthesis pathway and physiological data supported the collaborative relationship within the consortium that deserves further study.

## Acknowledgements

This study was supported by the Dutch Research Council through the UNLOCK project (NRGWI.obrug.2018.005), as well as Wageningen University through its Innovation Program Microbiology. We thank Peng Peng for giving expert guidance for isolation of anaerobes. We acknowledge the China Scholarship Council (CSC) for the support to Chen Zhang (File No. 201807720048).

## Data availability

The nucleotide sequence data has been deposited in the European Bioinformatics Institute under accession number PRJEB61380.

## Supplementary Information

**Figure S1.**
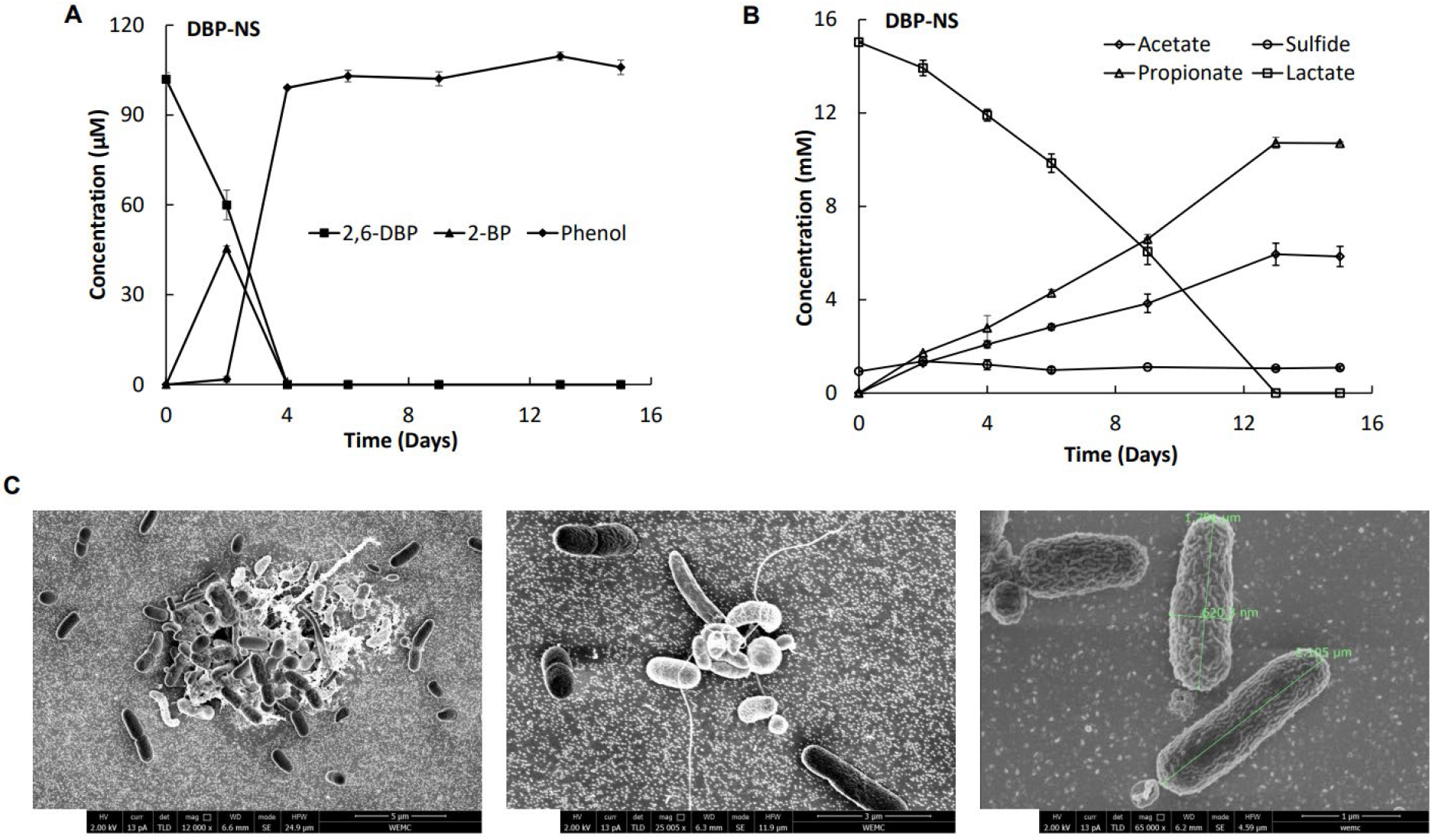
Reductive debromination of 2,6-dibromophenol (2,6-DBP) under sulfate-free conditions and scanning electron microscopy (SEM) of the consortium (related to Figure 1). 2,6-DBP was debrominated into phenol with the formation of 2-bromophenol as the intermediate (2-BP) (A). Lactate was consumed with the formation of propionate and acetate (B). The consortium was visualized by SEM at various magnifications, 12000 X (left), 25005 X (middle) and 65000 X (right) (C). The rod-shaped bacterium was measured in diameter (620.3 nm) and length (1.791 – 2.105 μm) as shown in green lines and letters.

**Figure S2.**
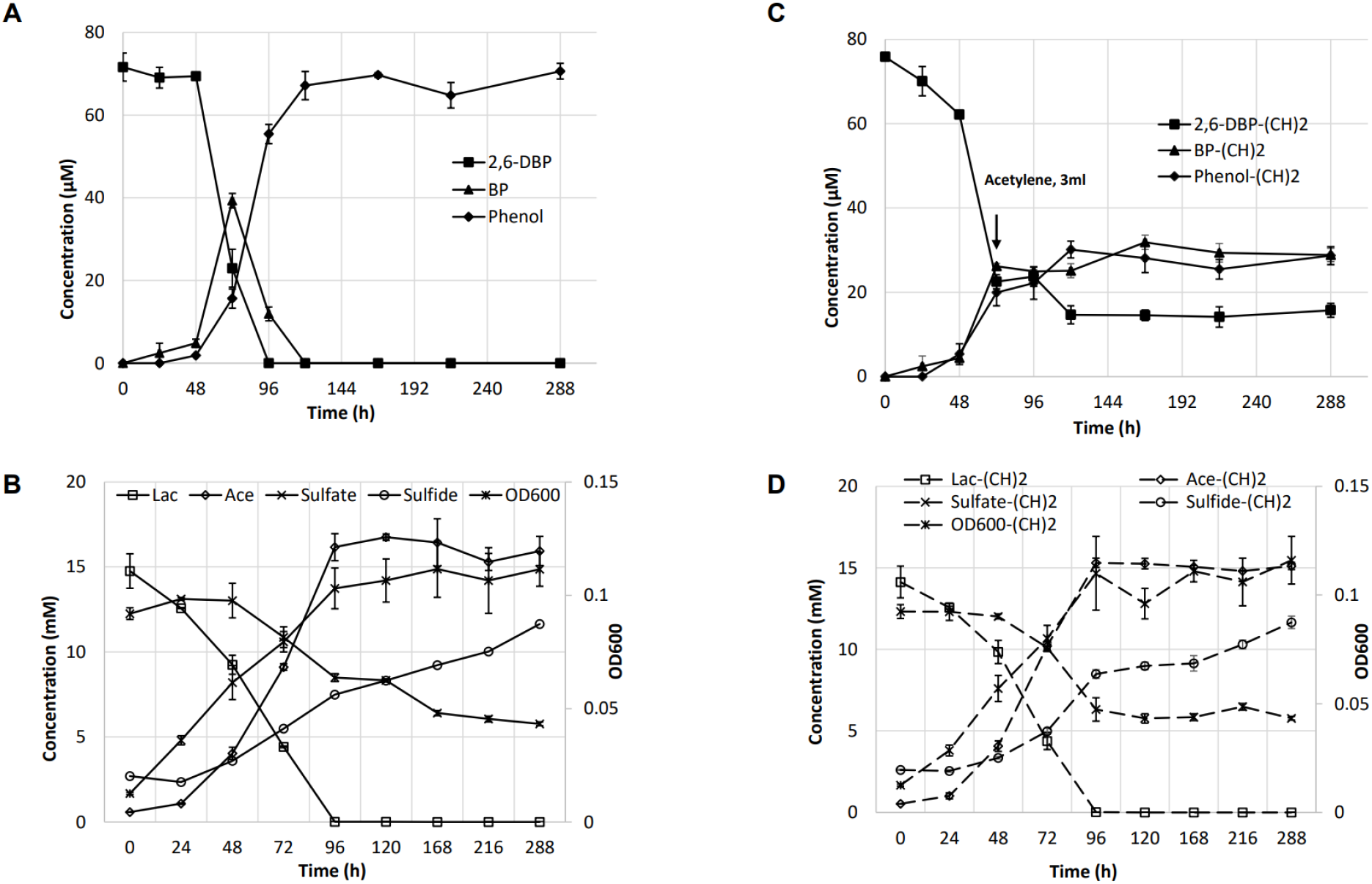
Inhibition by acetylene specifically on reductive debromination of 2,6-DBP in the presence of sulfate reduction (related to Figure 2). Reductive debromination of 2,6-DBP in addition to sulfate as the electron acceptor, and metabolite measurement with lactate as the electron donor and carbon source in the absence of acetylene (A, B), and presence of acetylene (C, D) respectively. (CH)2: acetylene; downward arrow indicates the injection of acetylene; Dashed lines represent the metabolism of lactate and sulfate after the addition of acetylene. Three replicate bottles were set, and the data indicate the mean ± SD. Error bars represent the SD.

**Figure S3.**
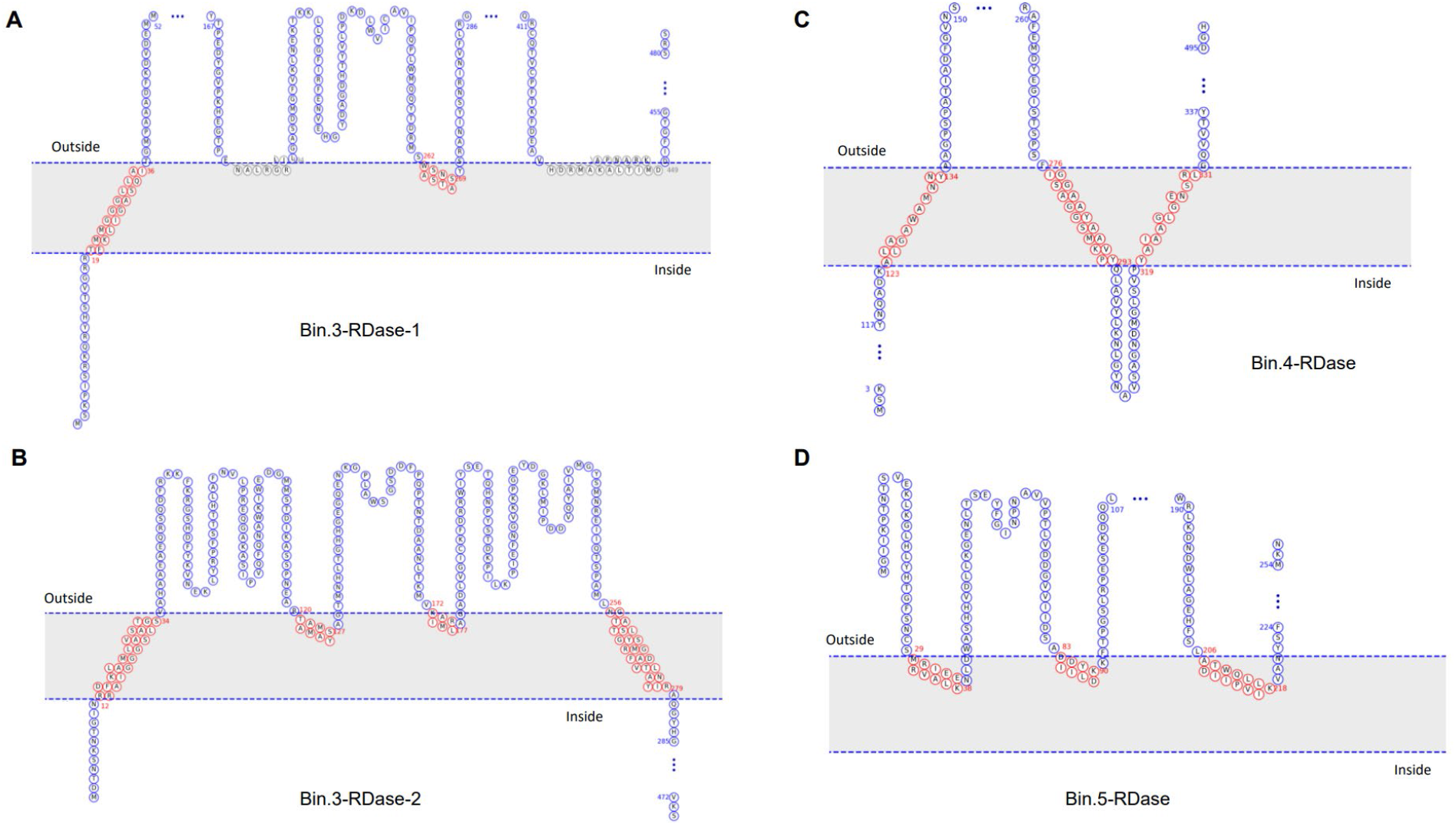
Transmembrane configuration prediction of RDases and Tph-RDase of assembled bins (related to Figure 4), including two RDases from bin.3 (A, B); one RDase from bin.4 (C) and TPh-RDase from bin.5 (D). The prediction of transmembrane configuration of the (TPh-) RDases was achieved by MemBrain (86).

**Figure S4.**
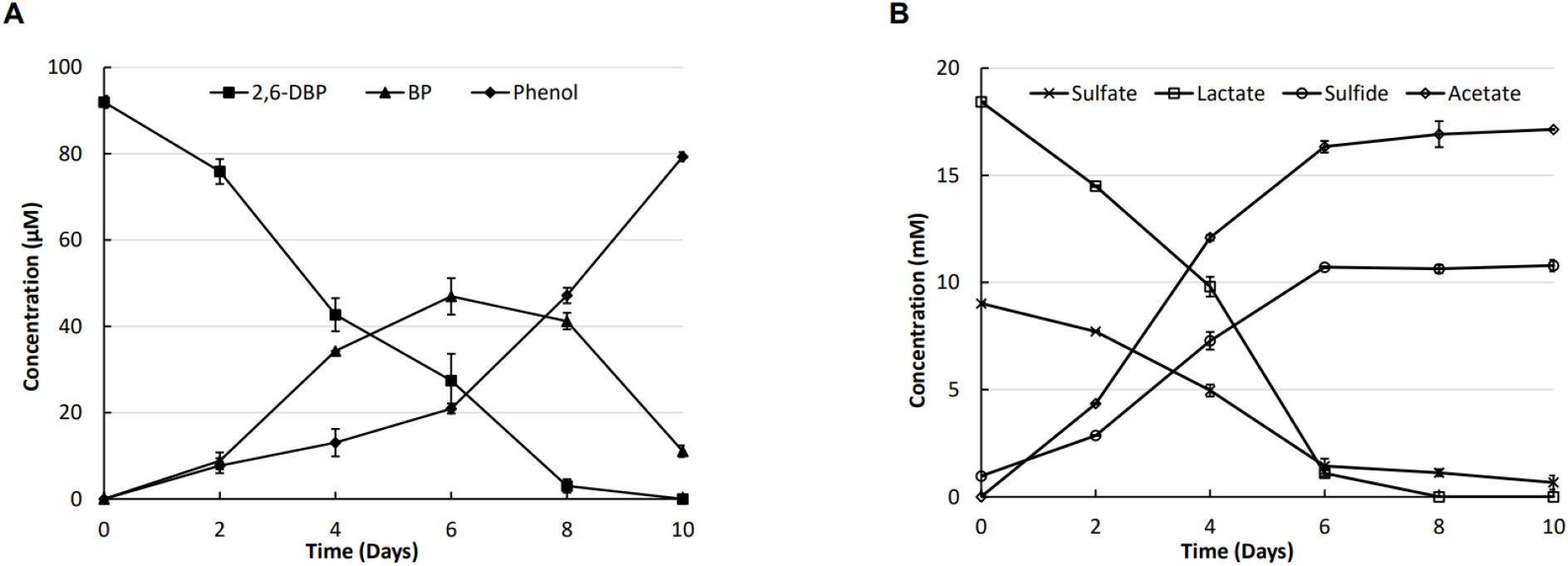
Reductive debromination of 2,6-DBP without additional B12 (related to Figure 5). Reductive debromination of 2,6-DBP to phenol with bromophenol as the intermediate (A). Lactate served as the electron donor and carbon source and was consumed to form acetate, and sulfate as the electron acceptor was reduced into sulfide (B). 2,6-DBP: 2,6-dibromophenol; BP: bromophenol; The experiment was set with three replicate bottles, and the data indicate the mean ± SD. Error bars represent the SD.

**Table S1.** Information of the assembled bins (MAGs)

| Bins | Classification | GC (%) | N50 (bp) | Size (bp) | Abundance | ANI<sup>a</sup> (%) |
| --- | --- | --- | --- | --- | --- | --- |
| bin.1 | *g_Oceanispirochaeta* | 43.1 | 10840 | 5074027 | 392.735242 | 93.85 |
| bin.2 | *g_Desulfoplanes* | 51.4 | 199123 | 3841128 | 1567.138705 | 78.11 |
| bin.3 | *g_Desulfoplanes* | 50.3 | 87237 | 4114339 | 1983.865769 | 78.14 |
| bin.4 | *f_Marinifilaceae* | 38.6 | 43215 | 5488154 | 523.39148 | N/A<sup>b</sup> |
| bin.5 | *g_Desulforhopalus* | 49.3 | 837424 | 3685696 | 9786.929752 | 76.45 |

<sup>a</sup> ANI: average nucleotide identity to the closest reference genome;
<sup>b</sup> N/A: indicates novel genus without closely related reference genome;

